# Direct lineage conversion of postnatal mouse cortical astrocytes to oligodendrocyte lineage cells

**DOI:** 10.1101/2024.05.28.596294

**Authors:** Justine Bajohr, Erica Y. Scott, Arman Olfat, Mehrshad Sadria, Kevin Lee, Maria Fahim, Hiba T. Taha, Daniela Lozano Casasbuenas, Ann Derham, Scott A. Yuzwa, Gary D. Bader, Maryam Faiz

## Abstract

Oligodendrocyte lineage cells (OLCs) are lost in many CNS diseases. Here, we investigate the generation of new OLCs via ectopic expression of *Sox10*, *Olig2* or *Nkxc.2* in mouse postnatal astrocytes. Using stringent analyses including, Aldh1l1-astrocyte fate mapping and live cell imaging we confirm that *Sox10* and *Olig2*, but not *Nkxc.2*, convert Aldh1l1^pos^ astrocytes to induced OLCs (iOLCs). With single cell RNA sequencing (scRNA-seq) we uncover the molecular signatures of iOLCs. Transcriptomic analysis of *Sox10*- and control cultures over time reveals a clear trajectory from astrocytes to iOLCs. Finally, perturbation models CellOracle and Fatecode support the idea that *Sox10* drives cells towards a terminal iOLC fate. Altogether, this multidimensional analysis shows bonafide conversion of astrocytes to iOLCs using *Sox10* or *Olig2* and provides a foundation for astrocyte DLR strategies to promote OLC repair.

## INTRODUCTION

Oligodendrocytes (OLs) are best known as the myelinating cells of the central nervous system (CNS). OLs ensheath neuronal axons to enable fast propagation of action potentials. Consequently, the loss or dysfunction of OLs results in impaired neurological function that is characteristic of many types of CNS disease and injury. Multiple sclerosis (MS), Alzheimer’s Disease, spinal cord injury, white matter stroke and cerebral palsy are all characterized by oligodendrocyte failure. Thus, therapeutic strategies aimed at replacing OLs are of significant clinical interest (1).

Direct lineage reprogramming (DLR) aims to generate new target cells lost to disease via the forced conversion of donor cells. Typically, DLR is performed by the overexpression of transcription factors (TFs). Early pioneering work from the Tesar Lab demonstrated fibroblast to induced oligodendrocyte progenitor cell (iOPC) conversion by ectopic expression of *Olig2*, *Sox10*, and *Nkxc.2*; determinants of OL cell fate in the embryonic brain (2). The combination of *Sox10* and *Olig2* was also later used to generate iOPCs from pericytes (3). Therapeutically, however, these newly generated iOPCs still require transplantation into the brain. Therefore, methods to reprogram endogenous CNS cells would be advantageous.

In parallel, astrocytes, CNS-resident cells, have emerged as donor cells in DLR strategies aimed at generating new neurons (4–7). Astrocytes are an attractive donor cell type for OL conversion given their shared neural origin (8,9). Astrocytes may already have relevant epigenetic marks and active TFs that could make DLR faster or more efficient (10–13). In addition, closely related cells may require fewer TFs for conversion. Indeed, conversion of astrocytes to an induced oligodendrocyte-like cell was suggested using *Sox10* alone (14). Therefore, we reasoned that single ‘Tesar’ factors, *Olig2*, *Sox10*, or *Nkxc.2,* could be used to force astrocyte conversion to new, induced oligodendrocyte lineage cells (iOLCs).

The OL lineage is comprised of oligodendrocyte progenitor cells (OPCs) that give rise to committed oligodendrocyte progenitors (COPs). These COPs differentiate into mature OLs (mOLs), comprising at least 6 different states (15). Of interest, during OL development, *Olig2, Sox10, and Nkxc.2* show different temporal expression and play different roles in OL fate specification. *Olig2,* long considered an OL lineage fate determinant (16), is also expressed in astrocytes (17,18), which suggests a broad role in early glial commitment (19,20). *Sox10* is important throughout OL development; *Sox10* promotes early OL lineage specification by re-inducing *Olig2* (21) and by inhibiting *Sufu (22)*, but is also required for OL survival following myelination (23,24). In contrast, *Nkxc.2* is expressed late in OL development, with myelin genes *Mbp* and *Mog,* and plays a role in regulating myelination (25,26). Given the unique roles of each of these TFs in development, we further hypothesized that each of these single factors would have varying efficiency in generating iOLCs.

Recent controversy in the field of astrocyte to neuron DLR *in vivo (27)* has highlighted the need for rigorous reporting of DLR outcomes. In a landmark study using astrocyte fate mapping strategies, the authors suggested that off-target transduction of endogenous cells was misrepresented as DLR (27). Therefore, it is important that new DLR paradigms utilize fate mapping, and stringent, multi-faceted analysis to determine the origin of the newly generated cells.

Here, we used lentiviral delivery of *Olig2*, *Sox10*, or *Nkxc.2* to investigate single TF conversion of postnatal (P0-P5) GFAP+ cortical astrocytes to iOLCs. Lineage tracing experiments using Aldh1l1-CreERT2;Ai14 mice confirmed that *Sox10* and *Olig2* directly convert Aldh1l1+ astrocytes to iOLCs. Moreover, live cell imaging, single cell RNA sequencing (scRNA seq) and deep learning methods support the findings that iOLCs can be generated from astrocytes following TF delivery. Altogether, these findings show bonafide astrocyte to iOLC DLR and lay the groundwork for future studies utilizing DLR for diseases involving OLC dysfunction and loss.

## METHODS

### Animals

All experiments were performed in accordance with approved Animal Use Protocols (AUP 20012151, 25-0389H) from the Division of Comparative Medicine at the University of Toronto. P0-P5 Ai14 (B6;129S6-*Gt(ROSA)2cSor^tm14(CAG-tdTomato)Hze^*/J, RRID:IMSR_JAX:007908) and Ai14;Ald1l1-CreERT2 mice (Ai14 crossed to B6N.FVB-Tg(Aldh1l1-cre/ERT2)1Khakh/J [RRID:IMSR_JAX:031008]) were used to generate postnatal astrocyte cultures.

### Cell Culture

Cortical astrocytes were isolated from male and female P0-P5 mice as previously described (28). Briefly, mice were decapitated, followed by the removal of the skull and meninges. Cortices were dissected, pooled, and mechanically dissociated in astrocyte media [DMEM (Gibco Catalogue No. 10569-010), 10% fetal bovine serum (FBS) (Gibco Catalogue No. 10082147) and 1% penicillin/streptomycin (Gibco Catalogue No. 15140122)]. Cells were cultured in flasks pre-coated with 10 µg/ml poly-d-lysine (Sigma Catalogue No. P6407), and incubated at 37°C, 5% CO2. Media was changed the day following isolation and every other day thereafter. Once the cells reached 80% confluency, typically after 6 days, flasks were placed on an orbital shaker for 30 minutes to remove contaminating microglia. Astrocyte media was replaced. For Ai14;Aldh1l1-CreERT2 cultures, 1uM 4-OHT was added to the astrocyte media at this step. Flasks were returned to the orbital shaker overnight, followed by vigorous shaking for one minute to remove contaminating OLCs. Media was removed and astrocytes were incubated in TrypLE Express Enzyme (Gibco Catalogue No. 12604013) for 5 minutes at 37°C, 5% CO2 to lift off the astrocytes. To inactivate the enzyme, astrocyte media was added at a 3:1 ratio (media:TrypLE). Cell suspension was collected and centrifuged at 300 x g for 5 minutes. Following removal of the supernatant, the pellet was resuspended in astrocyte media. Cells were then plated on poly-l-ornithine/laminin coated coverslips at either 50,000 or 70,000 cells/well in 24 well plates and incubated at 37°C, 5% CO2. For live cell analysis, cells were plated at 10,000 cells/ well in 96 well plates with poly-l-ornithine/laminin coating. For poly-l-ornithine/laminin coating, 0.1mg/ml poly-l-ornithine (Sigma Catalogue No. P4957) was added to dishes overnight at 37°C, washed 3 times with 1X phosphate buffered saline (PBS), and then dishes were incubated for two hours at 37°C with 10 μg/ml laminin (Sigma Catalogue No. L2020).

### Reprogramming

Lentiviral particles were purchased from VectorBuilder. For Ai14 reprogramming, LV-hGFAP::*Sox10-*P2A-Cre, LV-hGFAP::*Olig2*-P2A-Cre, LV-hGFAP::*Nkxc.2*-P2A-Cre, and control LV-hGFAP::BFP-T2A-Cre were used. For Ai14;Aldh1l1-CreERT2 reprogramming, LV-hGFAP::*Sox10*-P2A-zsGreen, LV-hGFAP::*Olig2*-P2A-zsGreen, LV-hGFAP::*Nkxc.2*-P2A-zsGreen and control LV-hGFAP::zsGreen were used. A multiplicity of infection (MOI) of 100 was used for all experiments. Virus-containing astrocyte media was placed on the cells and left overnight. Viral media was replaced with fresh astrocyte media one day post transduction (DPT). Three DPT, cells were switched to OPC differentiation media (2) [DMEM/F12/Glutamax (Gibco Catalogue No. 11330032), 1X N2 (Gibco Catalogue No. 17502048), 1X B27 without vitamin A (Gibco Catalogue No. 17504044), 200 ng/ml SHH (RCD Systems Catalogue No. 464-SH), 20 ng/ml FGF (RCD Systems Catalogue No. 3139-FB), 4 ng/ml PDGF (Sigma Catalogue No. SRP3228)]. At 10 DPT, the cells were switched to OL differentiation media (2) [DMEM/F12/Glutamax, 1X N2, 1X B27 without vitamin A, 40 ng/ml T3 (T2877 Catalogue No. Sigma), 200 ng/ml SHH, 100 ng/ml Noggin (RCD Systems Catalogue No. 1967-NG), 10 µM cAMP (Sigma Catalogue No. A9501), 100 ng/ml IGF (RCD Systems Catalogue No. 791-MG), 10 ng/ml NT3 (Sigma Catalogue No. SRP6007)].

### Live Cell Analysis

Astrocytes isolated from Ai14;Aldh1l1-CreERT2 mice were plated in 96 well plates, transduced and imaged every hour from 7 to 12DPT using the Apotome live cell system (Zeiss). 25 z-stack tiled images per well were captured with brightfield as well as the 488nm and 568nm fluorescent wavelength. Images were stitched to create a continuous video for each well. At 12DPT, cells were fixed and stained for OLC markers to confirm fate. Each well was then re-imaged at this final timepoint and OLC+ reprogrammed cells were matched to the live cell video and retrospectively analyzed for starting cell morphology and fluorescent expression. To characterize starting cell morphology, an analysis was first performed on 70 Aldh1l1+tdTomato+ astrocytes in tamoxifen-labelled Aldh1l1-CreERT2;Ai14 cultures and 70 PDGFRa+ OPCs from iOLC cultures (Table S1). Astrocyte and OPC paramenters were established for cell size, nucleus size, number of branches and branch thickness (Supplemental Figure 1). These parameters were then used to characterize starting cells as astrocytes or OPCs.

### Immunocytochemistry

Cells were fixed in 4% paraformaldehyde (PFA) (Sigma Catalogue No. P6148) for 20 minutes followed by three washes with 1X PBS. Cell membranes were permeabilized with 0.1% Triton-X-100 (Sigma Catalogue No. X100) for 10 minutes at room temperature, followed by three washes with 1X PBS, and then blocked with 5% milk for one hour at room temperature. Cells were incubated with primary antibodies in 1X PBS overnight at 4°C, washed three times with 1X PBS, and then incubated with secondary antibodies and DAPI (Sigma Catalogue No. D9542) in 1X PBS at room temperature for one hour. Following three final 1X PBS washes, coverslips were mounted on glass slides (Fisher Scientific Catalogue No. 125523) with Mowiol mounting solution (Sigma Catalogue No. 81381). For staining of membrane bound proteins (O4, PDGFRα), no permeabilization step with Triton-X-100 was performed. Primary antibodies: mouse anti-SOX10 (RRID:AB_10844002, 1:250), rabbit anti-PDGFRα (RRID:AB_2892065, 1:500), mouse anti-O4 (RRID:AB_357617, 1:1000) and rat anti-MBP RRID:AB_305869, 1:50).

Secondary antibodies: anti-mouse IgG 488 (Invitrogen Catalogue No. A32723) and 647 (Invitrogen Catalogue No. A32728), anti-mouse IgM heavy chain 488 (Invitrogen Catalogue No. A21042) and 647 (Invitrogen Catalogue No. A21238), anti-rabbit 488 IgG (Invitrogen Catalogue No. A11034) and 647 (Invitrogen Catalogue No. A32733), anti-rat IgG 488 (Invitrogen Catalogue No. A21208) and 647 (Invitrogen Catalogue No. A21247) all at 1:1000.

### Microscopy and Image Analysis

Fluorescent images for quantification were taken on an LSM 880 Elyra Superresolution and LSM 900 (Zeiss) using a 20x objective and Zen Blue software (Zeiss). Post-acquisition linear adjustments of brightness for all channels were made to micrographs using the Zen Blue software in Figure 1C, Figure 2E-G and Supplemental Figure 5D. For quantification, ten regions of interest were selected at random for each well. Images were analyzed using ImageJ software (National Institutes of Health, RRID:SCR_003070). Reprogramming efficiency was calculated as a measure of total OLC marker^+^reporter^+^DAPI^+^ cells over total reporter^+^DAPI^+^cells.

**Figure 1.**
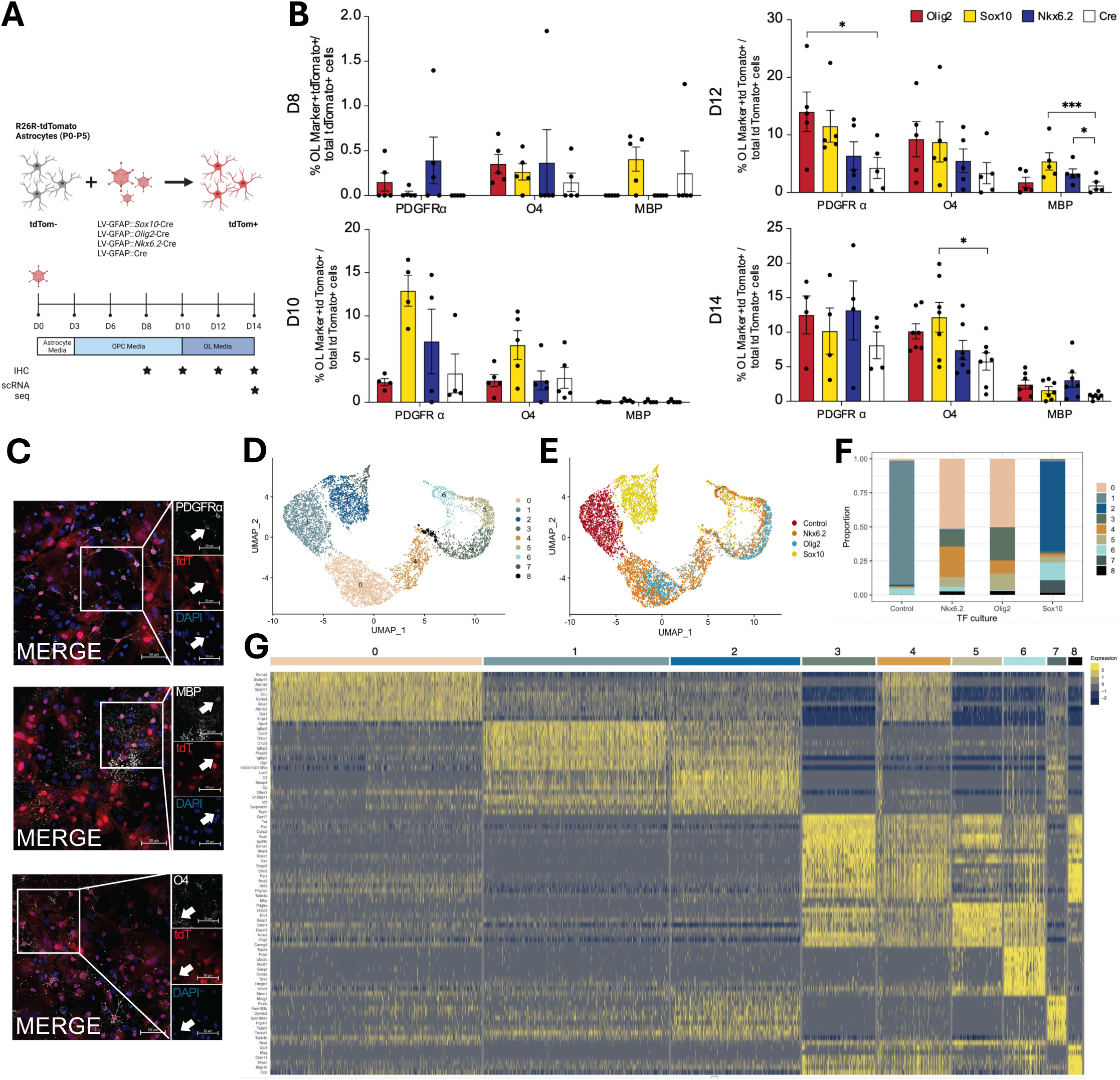
*Sox10*, *Olig2* and *Nkxc.2* convert GFAP+ cells to oligodendrocyte lineage cells. **(A)** Experimental design and timeline. **(B)** Quantification of PDGFRα^+^tdTomato^+^iOPCs, O4^+^tdTomato^+^ iCOPs and MBP^+^tdTomato^+^ iOLs at 8 (n=5), 10 (n=4), 12 (n=5) and 14 (n=7) days post transduction (DPT). Data are presented as mean ± SEM, each data point represents one individual cell culture experiment; at each time point and for each cell type marker, a matched pairs one-way ANOVA or Kruskal-Wallis (d8, d10, d12) or one-way ANOVA (d14) was performed with Dunnet’s post testing (*= p<0.05, *** = p< 0.001). **(C)** Representative images of PDGFRα^+^tdTomato^+^,O4^+^tdTomato^+^ and MBP^+^tdTomato^+^ cells at 12DPT. Single channel images are shown of the boxed cells (arrows indicate double positive cells). Scale bar = 50um. **(D)** UMAP clustering of *Olig2*-, *Sox10*-, *Nkxc.2*- and control (Cre) transduced cells at 14DPT **(E)** UMAP of (D) overlayed with treatment (*Olig2*-, *Sox10*-, *Nkxc.2*- and control (Cre)) **(F)** Proportion analysis of clusters found in *Olig2*-, *Sox10*-, *Nkxc.2*-, and control (Cre) transduced cells. **(G)** Heatmap of top upregulated genes from each cluster in (D).

**Figure 2.**
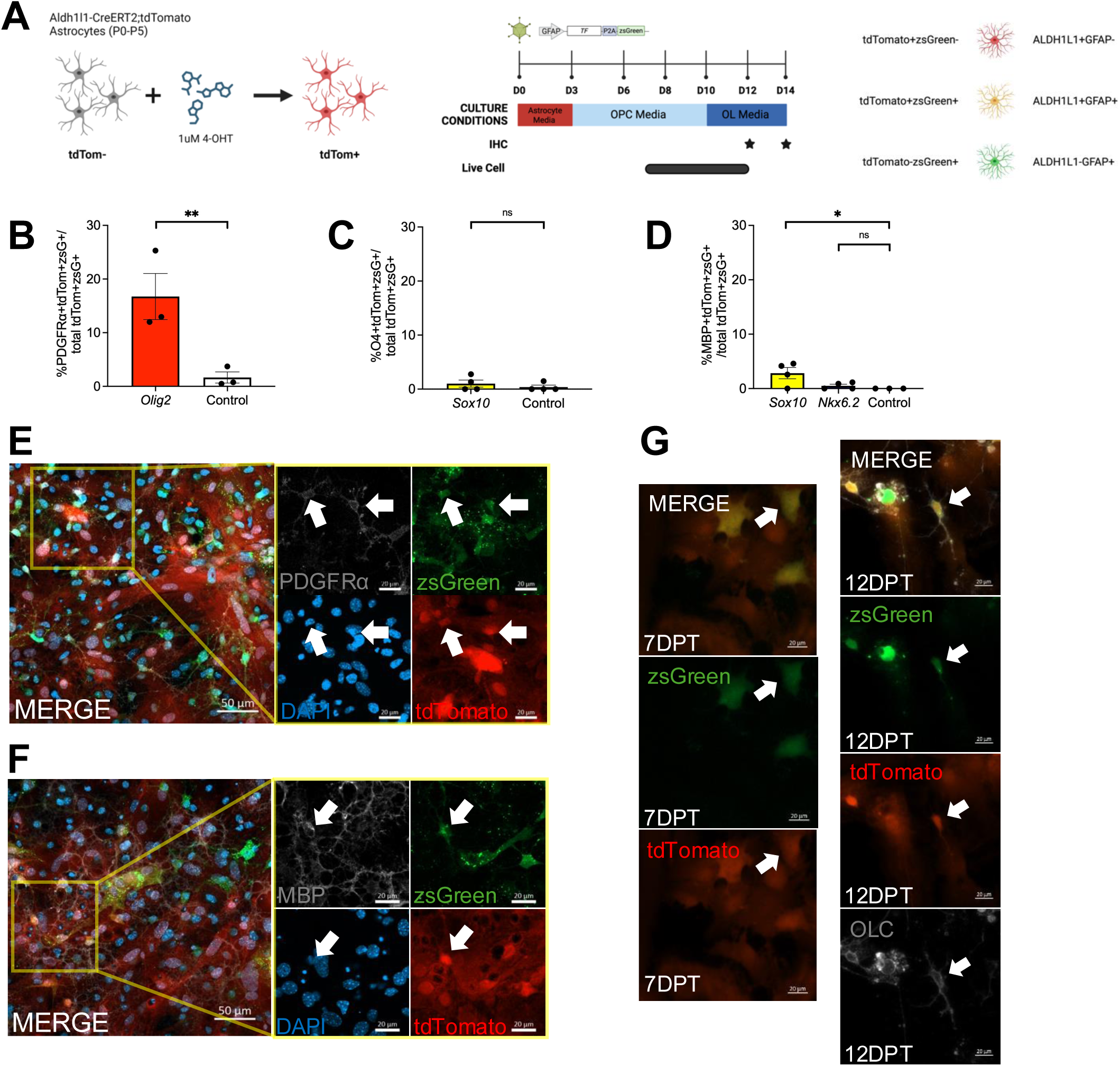
Lineage tracing confirms true conversion of astrocytes to oligodendrocyte lineage cells. **(A)** Experimental design, timeline and outcomes. **(B)** Quantification of tdTomato^+^zsGreen^+^PDGFRα^+^ iOPCs at 12DPT (n= 3). Data are presented as mean ± SEM, each data point represents one individual cell culture, a paired t-test was performed (** = p<0.01). **(C)** Quantification of tdTomato^+^zsGreen^+^O4^+^ iCOPs at 14DPT (n = 4). Data are presented as mean ± SEM, each data point represents one individual cell culture, a Wilcoxon test was performed (ns). **(D)** Quantification of tdTomato^+^zsGreen^+^MBP^+^ iOLs at 12DPT (n= 4 for *Sox10, Nkxc.2*, n= 3 for Control). Data are presented as mean ± SEM, each data point represents one individual cell culture, a one-way ANOVA with Dunnet’s post testing was performed (* = p<0.05). **(E)** Representative image of PDGFRα^+^tdTomato^+^zsGreen^+^ cells 12DPT. Single channel images are shown of the boxed cells (arrows indicate triple positive cells, scale bar = 50um (merge) and 20um (single channel)). **(F)** Representative image of MBP^+^tdTomato^+^zsGreen^+^ cells 12DPT. Single channel images are shown of the boxed cells (arrows indicate triple positive cells, scale bar = 50um (merge) and 20um (single channel)). **(G)** Representative tdTomato^+^zsGreen^+^ cell with astrocyte-like morphology at onset (7DPT) and OLC expression at the end (12DPT) of live cell tracking (arrow indicates tracked cell, scale bar = 20um).

### scRNA-seq capture and processing

At 14DPT, LV-hGFAP::*Sox10-*P2A-Cre, LV-hGFAP::*Olig2*-P2A-Cre, LV-hGFAP::*Nkxc.2*-P2A-Cre, and control LV-hGFAP::BFP-T2A-Cre cultures were processed using the BD Rhapsody System (BD Biosciences) and then sequenced. For single-cell isolation, an average of 9813.75 viable cells were captured in wells at cell load (Table S2). The BD Rhapsody scanner reported an average multiplet rate of 10.13% and an average number of wells with viable cells and a bead of 7081.5 (Table S2). Detailed metrics for each sample can be found in Table S2. Samples were down-sampled to 2500 cells and carried through and converted to cDNA using the BD Rhapsody WTA Reagent Kit (Becton Dickinson Canada, Catalogue No. 633802). Each cell was sequenced at approximately 100 million reads per cell (at least 2x150 bp paired-end reads) on a Novaseq (Donnelly Sequencing Centre, University of Toronto).

In addition, LV-GFAP::*Sox10* and control LV-GFAP::Cre cultures were collected prior to transduction, at 3DPT and 8DPT, and processed using the BD Rhapsody System (BD Biosciences) and then sequenced. For single-cell isolation, an average of 10263.8 viable cells were captured in wells at cell load (Table S3). The BD Rhapsody scanner reported an average multiplet rate of 6.86% and an average number of wells with viable cells and a bead of 7785 (Table S3). Detailed metrics for each sample can be found in Table S3. Samples were down-sampled to 2500 cells and carried through and converted to cDNA using the BD Rhapsody WTA Reagent Kit (Becton Dickinson Canada, Catalogue No. 633802). Each cell was sequenced at approximately 100 million reads per cell (at least 2x150 bp paired-end reads) on a Novaseq (Donnelly Sequencing Centre, University of Toronto).

### scRNA-seq analysis

Fastq files were first demultiplexed with Kallisto (29) (RRID:SCR_016582)(v0.48.0) and Bustools (30) (RRID:SCR_018210)(V 0.41.0) using supplied whitelists (Data S1) with the - BDWTA option and aligning to GRCm38.96 with Cre sequence appended to the end. Bustools (30) was then used to generate gene count tables. Cells were plotted based upon UMI counts per barcode, thresholds were selected based on inflection point of UMI count per barcode plots. These thresholds produced read and gene count distributions that were comparable between all treatment groups (Supplemental Figure 2A, Supplemental Figure 6A). Gene count tables were made into S4 objects, scaled, normalized and dimensions reduced (PCA then UMAP) using the Seurat (31) package (RRID:SCR_016341) (v4.1). Clusters identified as microglia and as vascular and leptomeningeal cells (VLMCs) (Supplemental Figure 2B-C [cluster 8], Supplemental Figure 6B-C [cluster 6 and 8]) were removed prior to further analysis.

Gene markers for oligodendrocyte lineage cells were adopted from studies observing *in vivo* mouse oligodendrocyte lineage cells across several areas in young and mature CNS tissues (15). These markers were converted into percent expression of each UMAP cluster using the PercentageFeatureSet function from Seurat (31), with further gene resolution displayed by heatmaps created using ComplexHeatmap (32)(RRID:SCR_017270) (v12.13.1). An additional set of gene markers, demonstrating similar, but less resolved, conclusions was also used from an *in vitro* rat study looking at OLCs from the cortex (33). Stacked violin cluster dot plots were made with scCustomize (34)(RRID:SCR_024675)(v2.1.2). For pseudotime and trajectory analysis Slingshot (35) (RRID:SCR_017012)(v2.3.1) and Monocle3 (36–40) (RRID:SCR_018685) were used. For *in silico Sox10* perturbation, CellOracle (41) (v0.14.0) was used. Plots were produced using ggplot2 (42) (RRID:SCR_014601)(v3.3.5) and figure generating scripts were run in R studio (43) (v4.2.0), with demultiplexing using Kallisto (29) and Bustools (30) run on a Compute Canada HPC cluster. All scripts used for processing of scRNA-seq data and for figure generation can be found at github.com/eyscott/Bajor_BDRhapsodyData_DLR.

### Deep-learning analysis

Fatecode (44) was used to identify key genes for cellular transition. To optimize the autoencoder and subsequent classifier configuration, a grid search was conducted to systematically evaluate various combinations of hyperparameters. These included: latent layer size, number of nodes in the first and second layers of the autoencoder, classifier architecture, and type of activation function. The grid search aimed to identify the hyperparameters that minimized a combined reconstruction and classification loss function, signifying the optimal performance for our specific dataset. Perturbations on the latent space were performed and cell classifications as well as genes associated with each perturbation were obtained.

### Statistical Analysis

Percentage values were transformed using the arcsine square root transformation and assessed for normal distribution using the Shapiro-Wilks test. When distribution and variance were equal, a matched pairs one-way ANOVA ([Ai14:D8, D10, D12 O4, D12 MBP], [Ai14;Aldh1l1-CreERT2: D12 MBP zsGreen+, D12 MBP tdTomato+) or one-way ANOVA ([Ai14:D14], [Ai14;Aldh1l1-CreERT2: D12 MBP zsGreen+,tdTomato+]), or paired t-test ([Ai14;Aldh1l1-CreERT2: D12 PDGFRα zsGreen+tdTomato+, D12 PDGFRα tdTomato+, O4 zsGreen+, O4 tdTomato+]) was performed to compare reprogramming efficiency of TF groups to a control group (Ai14: LV-GFAP::Cre, Ai14;Aldh1l1-CreERT2: LV-GFAP::zsGreen). When transformed values did not follow a Gaussian distribution, a Kruskal-Wallis test (Ai14: D12 PDGFRα, D14 PDGFRα) or Wilcoxon test (Ai14;Aldh1l1-CreERT2: PDGFRα zsGreen+, O4 zsGreen+tdTomato+) was performed to compare reprogramming efficiency to a control group (Ai14: LV-GFAP::Cre, Ai14;Aldh1l1-CreERT2: LV-GFAP::zsGreen). In both cases, Dunnett’s post-hoc testing was performed to correct for multiple comparisons. Differences were considered significant at p < 0.05. Values are presented as mean ± SEM. The statistical software used for transformation, distribution, variance, ANOVA, Kruskal-Wallis, t-test, Wilcoxon test and Dunnett’s analysis was GraphPad Prism version 9.0.1 (RRID:SCR_002798).

## RESULTS

### iOLCs are generated following expression of *Olig2*, *Sox10* or *Nkxc.2*

To investigate astrocyte to OLC conversion, we established postnatal, cortical astrocyte cultures from Ai14 mice. To understand the purity of our cultures, we quantified the numbers of contaminating OLCs. Quantification of SOX10+, O4+ and MBP+ OLCs showed less than 2.5% in our cultures (Supplemental Figure 3A).

To examine the reprogramming potential of *Olig2, Sox10,* and *Nkxc.2*, we transduced Ai14 astrocytes with either LV-GFAP::*Olig2,* LV-GFAP::*Sox10*, LV-GFAP::*Nkxc.2* or a control LV-GFAP::*Cre* (Figure 1A). When we quantified the numbers of tdTomato+ cells that co-expressed OPC (PDGFRa), COP (O4) or OL (MBP) markers at 8 and 10 DPT, no differences were seen in any of the TF-transduced cultures compared to controls (Figure 1B). By 12 DPT, *Olig2*-cultures showed an increase in the percentage of tdTomato^+^PDGFRα^+^ OPCs (p = 0.0309, H-statistic: 7.514), whereas *Sox10*- and *Nkxc.2*-cultures showed an increase in tdTomato^+^MBP^+^ OLs (*Sox10:* p = 0.0004, *Nkxc.2:* p = 0.0108, F statistic = 13.28, degrees of freedom =3) (Figure 1B-C). No differences were seen in the percentage of tdTomato^+^O4^+^ COPs in any condition at 12DPT (Figure 1B). However, by 14DPT we observed an increase in the percentage of tdTomato^+^O4^+^ COPs in *Sox10-* cultures when compared to controls (p = 0.0213, F statistic = 3.512, degrees of freedom = 3) (Figure 1B-C). Finally, to understand whether a longer time in culture could increase the number of MBP+ OLs, we cultured cells for an additional 8 days and analyzed the percent of tdTomato^+^MBP^+^ cells at 22 DPT. No differences were seen in tdTomato^+^MBP^+^ OLs in any condition compared to controls (Supplemental Figure 3B). Altogether, these findings suggest that *Olig2*, *Sox10* and *Nkxc.2* increase the percentage of iOLCs relative to controls in Ai14 astrocyte cultures.

### Canonical OLC cluster found in TF-treated cells following scRNA-seq

To further characterize *Olig2-, Sox10-, Nkxc.2-* and control cultures, we performed scRNA-seq at 14DPT (Figure 1A). Clustering analysis showed the appearance of nine clusters (Figure 1D). We then used proportion analysis to identify clusters that were unique to TF-treated samples. Clusters 3 and 8 were predominantly comprised of cells from TF-induced cultures (Figure 1E-F). The top 10 genes marking each cluster in Figure 1D are highlighted in Figure 1G. Canonical OLC genes such as *Mbp, Plp1, Bcas1* and *Cldn11* were expressed in clusters 3 and 8 (Figure 1G), further suggesting that the increase in iOLCs is a result of TF delivery.

Next, we compared the molecular profiles of clusters 3 and 8 to established OL lineage datasets. First, we used OLC specific annotations of mouse fetal and adult OLCs derived from Marques *et al* (15) to bin the data. We found that cluster 3 was characterized by COP signatures, while cluster 8 was characterized by COP, newly formed oligodendrocyte (nfOL) and myelin forming oligodendrocyte (mfOL) signatures (Supplemental Figure 4A). We also used OLC annotations of *in vitro* OLCs derived from Dugas *et al* (33) to bin the data. Again, we saw that cluster 8 was characterized by OL signatures, but cluster 3 was not. (Supplemental Figure 4B). Altogether, these findings further support the observation of increased OLC generation in TF-induced Ai14 cultures (Figure 1B).

### Bonafide astrocyte to OLC conversion is confirmed with astrocyte fate mapping

We had previously observed contaminating OLCs in our cultures and the presence of tdTomato^+^OLC marker^+^ cells in our Cre controls. Therefore, to confirm the origin of these newly generated OLCs, we performed astrocyte fate mapping using Ai14;Aldh1l1-CreERT2 cultures, the current gold standard in the field (27). Prior to transduction, post-natal cortical Ai14;Aldh1l1-CreERT2 astrocyte cultures were treated with 1uM 4-OHT to permanently label all *Aldh1l1* expressing cells with tdTomato (Figure 2A). Cultures were transduced with LV-GFAP::*Sox10*-zsGreen, LV-GFAP::*Olig2*-zsGreen, LV-GFAP::*Nkxc.2*-zsGreen or a LV-GFAP::zsGreen control (Figure 2A). Any OLCs that were directly converted by a TF from these *Aldh1l1* expressing astrocytes would therefore express zsGreen and tdTomato (Figure 2A). To understand whether the OPCs generated from *Olig2* overexpression were the product of astrocyte reprogramming, we quantified the percentage of PDGFRα+zsGreen^+^tdTomato^+^ cells at 12DPT. We observed an increase in PDGFRα^+^zsGreen^+^tdTomato^+^ OPCs compared to controls, confirming the astrocytic origin of iOPCs (Figure 2B,E). To assess the origin of *Sox10*-induced COPs we quantified the percentage of O4^+^zsGreen^+^tdTomato^+^ cells at 14DPT. No increase was observed in *Sox10* cultures compared to controls (Figure 2C). Finally, to determine the origin of OLs generated by *Sox10* or *Nkxc.2,* we quantified the percentage of MBP^+^zsGreen^+^tdTomato^+^ OLs at 12DPT. We observed an increase of MBP^+^zsGreen^+^tdTomato^+^ OLs generated by *Sox10*, but not from *Nkxc.2* (Figure 2D,F). This suggests that *Olig2* and *Sox10* reprogram *Aldh1l1* astrocytes to iOLCs, while *Nkxc.2*-derived OLs and *Sox10*-derived COPs previously observed (Figure 1B), are not the result of *Aldh1l1* astrocyte conversion. To further confirm these findings, we performed live cell imaging of Ai14;Aldh1l1-CreERT2 *Sox10*-treated and control cultures from 7DPT to 12DPT. At 12DPT, we observed tdTomato^+^zsGreen^+^ cells that co-expressed OLC markers (PDGFRa/O4/MBP+) and showed OPC-like morphologies (small nucleus, multiple thin branches) (Figure 2G, Supplemental Figure 1). When we tracked these cells back to 7DPT, we found that they showed a characteristic astrocyte morphology (large cell body and nucleus, with minimal branching) (Figure 2G, Supplemental Figure 1, Supplemental Video 1). Taken together, these findings further demonstrate bonafide astrocyte to iOLC conversion.

We also examined the numbers of Aldh1l1^+^ non-transduced (tdTomato^+^only) cells, as well as Aldh1l1^neg^ virally transduced (zsGreen^+^ only) cells (Supplemental Figure 5). No difference in the percentage of OLC marker^+^tdTomato^+^zsGreen^neg^ was seen between TF-treated and control cultures (Supplemental Figure 5). However, when we examined zsGreen^+^tdTomato^neg^ cells, we observed an interesting increase in the percentage of MBP^+^zsGreen^+^tdTomato^neg^ OLs at 12DPT in *Sox10*-treated cells compared to controls (Supplemental Figure 5C). When we analyzed our live cell imaging experiment to understand the origin of these cells, we found examples of tdTomato^neg^zsGreen^+^OLCmarker^+^ cells that arose from a tdTomato^neg^ cell (Supplemental Figure 5D). These tdTomato^neg^ cells did not have OPC morphology 7DPT (Supplemental Figure 1,5D, Supplemental Video 2). Rather, they showed morphologies typical of astrocytes (Supplemental Figure 1,5D, Supplemental Video 2), suggesting the conversion of an astrocyte-like Aldh1l1^neg^ cell.

### Characterization of *Sox10*-mediated DLR using scRNA-seq

Given that *Sox10* increased the number of mOLs from *Aldh1l1* astrocytes (Figure 2D,F), we next probed the mechanisms underlying *Sox10*-mediated reprogramming. We performed scRNA-sequencing at three additional timepoints (D0 prior to transduction, 3DPT, and 8DPT) on Cre-control and *Sox10*-treated cultures. For analysis, we combined cells from D0, 3DPT, 8DPT and 14DPT (Supplemental Figure 6B). We excluded cells with microglia and VLMC markers (Supplemental Figure 6C). Clustering analysis of the remaining cells showed the appearance of 10 distinct clusters (Figure 3A). We characterized these clusters based on expression of genes associated with astrocytes, oligodendrocytes, and NG2 cells, as well as proliferation markers (Figure 3B-C).

**Figure 3.**
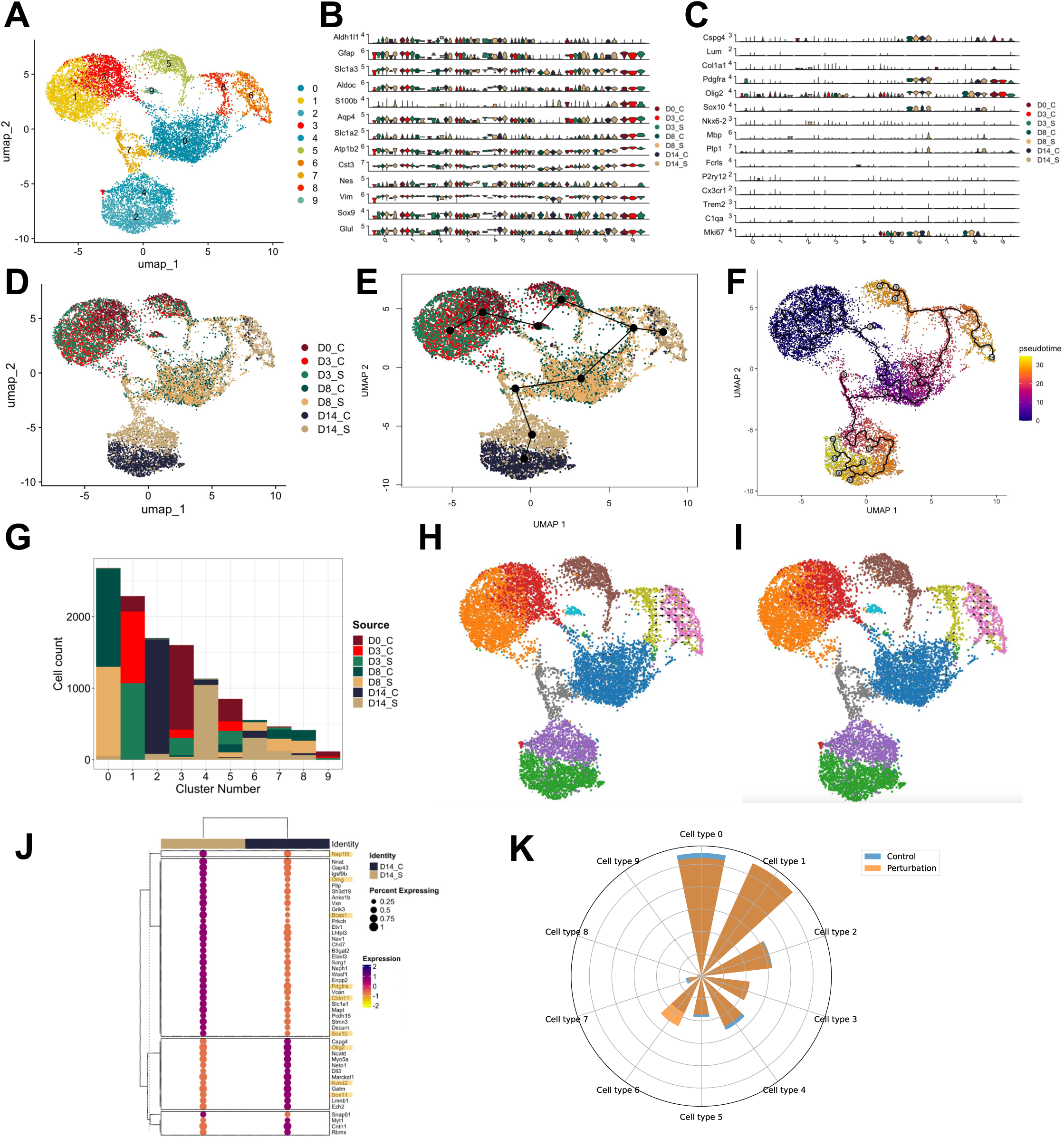
Characterization of DLR using scRNA-seq shows terminal oligodendrocyte cluster of cells at day 14 driven by *Sox10*. **(A)** UMAP clustering of *Sox10* and *Cre* control treated cells from prior to transduction, as well as 3, 8 and 14DPT **(B)** Canonical astrocyte gene expression (log-normalized, y-axis) separated by cluster (x-axis) and coloured by timepoint and treatment group. **(C)** Canonical gene expression (log-normalized, y-axis) of NG2 glia, VLMC, OLC, microglia and proliferation, separated by cluster (x-axis) and coloured by timepoint and treatment group. **(D)** UMAP clustering from (A) overlayed with timepoint and treatment group **(E)** Slingshot lineage analysis of *Sox10* and *Cre* control treated clusters overlaid with UMAP embeddings. **(F)** Monocle3 lineage analysis of *Sox10* and *Cre* control treated clusters overlaid on UMAP plot and coloured by pseudotime predictions. **(G)** The number of cells (y-axis) per cluster (x-axis) originating from each of the coloured timepoint+treatmentgroups. **(H)** CellOracle modeling of *in silico Sox10* knock out (KO) overlaid onto UMAP plot. Arrows indicate trajectory prediction with *Sox10* KO. **(I)** CellOracle modeling of *in silico Sox10* knock in (KI) overlaid onto UMAP plot. Arrows indicate trajectory prediction with *Sox10* KI **(J)** Clustered, top differentially expressed genes (dots, y-axis) between 14DPT *Sox10 (D14_S,* Beige, left) and *Cre (D14_C,* Blue, right) control treated cells from cluster 6. Size of dot is scaled to percent of cells in the cluster expressing that gene, colour of dot represents the average scaled expression of the gene across cells. **(K)** Predicted cell types following Fatecode perturbation on node 16 of the latent layer in the *Sox10* and control treated dataset. Blue represents the number of cells prior to perturbation. Orange (overlayed) represents the number of cells following perturbation.

First, we examined the molecular identities of cells in our starting cultures. D0 cells were found in clusters 1, 3, 5 and 9 and showed expression of canonical astrocyte markers, including *Slc1a3, Aqp4, Atp1b2, Glul* and *Gfap* (Figure 3B,D). Of interest, clusters 1, 3 and 5 showed expression of *Aldh1l1*, but this gene was absent in cluster 9 (Figure 3B), which could explain the conversion of tdTomato^neg^ cells observed in our fate mapping experiments (Supplemental Figure 5). Altogether, this suggests that early postnatal cultures are heterogeneous and comprised of distinct astrocyte-like populations (clusters).

To understand the cell fate transitions that occur over the reprogramming timecourse, we performed trajectory analysis with Slingshot (35) (Figure 3E) and Monocle3 (36–40) (Figure 3F). Slingshot and Monocle3 both reconstructed trajectories with clusters 6 and 4 as terminal branches (Figure 3E-F). We then examined the origin of cells comprising each cluster. As expected, the clusters formulating earlier roots of the Monocle3 branches (clusters 0, 1, 3, 9) were primarily comprised of D0, 3DPT, and 8DPT timepoints (Figure 3G). Clusters 6 and 4, were predominantly comprised of 14DPT samples, and in particular, cells from *Sox10*-treated samples (D14_S) (Figure 3G). Analysis of the differentially expressed genes (DEGs) at these branches showed that cluster 6 and branch 14 of the Monocle3 trajectory were enriched in the OLC genes *Sox10* (21–24) (p- and q-value=0), *Bcas1* (45) (p- and q-value=0), and *Omg* (46,47) (p- and q-value=0 (Supplemental Figure 7A). In contrast, cluster 4 and the Monocle trajectory branch 11 showed expression of the genes *Lcn2* (p- and q-value=0) and *Igfbp5* (p- and q-value=0), (Supplemental Figure 7B) previously shown to be expressed in reactive astrocytes (48–50). All Monocle3-derived differentially expressed genes can be found in Data S2. Altogether, these analyses suggest that the trajectory to branch 14/cluster 6 represents the path of astrocyte to OLC conversion.

To determine how *Sox10* would influence the gene regulatory network of cells in clusters 6 and 4, we performed *in silico* perturbation of *Sox10* using CellOracle (41). *in silico* knock out (KO) of *Sox10* predicted a large shift away from a cluster 6 identity and little change away from the identity of cluster 4 (Figure 3H). In agreement, *in silico* knock in (KI) of *Sox10* showed a large shift towards a cluster 6 identify and little change towards a cluster 4 identity (Figure 3I). This suggests that leaf 14 and cluster 6 represent cells with a gene regulatory network most affected by *Sox10*.

As cluster 6 was comprised of both *Sox10-*treated and Cre-control cells, we then performed a differential gene expression analysis of 14DPT Cre-control and *Sox10-*treated cells within this cluster. 175 DEGs were found, with the top genes (expressed in at least 50% of Cluster 6, *Sox10*-treated cells and with a difference of at least 0.5 pct between Cre-control and *Sox10*-treated cultures) clustered and provided in a dotplot (Figure 3J). This analysis highlighted that genes including *Omg*, *Bcas1*, *Pdgfra*, *Cldn11 (51)*, and *Sox10* are enriched in *Sox10*-treated cells compared to Cre-control cells, suggesting that *Sox10*-treated cells are more representative of OLCs than the Cre-control cells exposed to OPC and OL media alone.

### Understanding the genetic drivers of astrocyte to OLC DLR

Our analysis showed that *Sox10* was important for determining the OLC identity of cluster 6. We specifically chose *Sox10* based on its role in OLC fate specification and development (21–24). To understand whether there might be other [better] candidate genes that would promote an OLC identity, we used an unbiased, deep learning perturbation model, Fatecode (44), to predict genes that would allow cells to shift from a cluster 4 identity (astrocytes at 14DPT, not fully reprogrammed) to a cluster 6 identity (OLCs at 14DPT, end state of DLR). Following training of the Fatecode model on our *Sox10* and Cre-control treated DLR dataset, we identified perturbation on node 16 of the latent layer as one which increased the number of cells in cluster 6 whilst simultaneously reducing the number of cells in clusters 0, 2, 4, 5 and 7. (Figure 3K). Given that the total number of cells in the dataset does not change, this suggests that the perturbation on node 16 of the latent layer was pushing cells that did not reprogram (astrocytes at 8 and 14DPT) towards a reprogrammed (OLCs at 14DPT) fate. To identify the genes driving this shift,we ranked the absolute value of gene expression changes to obtain a list of 3000 genes involved in this perturbation, with the top 40 genes listed in Table S4. Strikingly, we observed that *Sox10* was ranked as the ninth most correlated gene and the top TF driving this shift (Table S4). This supports our previous findings that suggest *Sox10* drives astrocyte to iOLC DLR. Additionally, we observed genes important for myelination (*Plp1, Mbp*) (52), as well as other TFs previously identified with OLC differentiation and myelination (*Sox3* (53)*, KlfS* (54)) as integral to this shift (Table S4). Taken together, these findings further support the importance of *Sox10* in reprogramming astrocytes to iOLCs and identify additional genes and TFs that may be ideal candidates for astrocyte to iOLC reprogramming.

## DISCUSSION

Here, we show astrocyte to iOLC conversion with *Sox10* or *Olig2* using a battery of experimental tools, including astrocyte fate mapping, live cell imaging, scRNA-seq timecourse and unbiased deep learning. While previous studies demonstrated the generation of iOLCs from different types of somatic cells using combinations of *Sox10*, *Olig2* and *Nkxc.2* (2,3,55), ours is the first to compare the individual reprogramming ability of each these TFs in astrocytes. Using *Aldh1l1*-based fate-mapping, we found that that *Sox10* and *Olig2* convert Aldh1l1^+^ astrocytes to iOLCs.

In contrast to *Sox10* and *Olig2*, *Nkxc.2* was unable to convert *Aldh1l1^+^*astrocytes to new iOLCs. Unlike *Sox10* and *Olig2*, which define oligodendrocyte identity and continue to shape their gene regulatory network throughout life (56), *Nkxc.2* may be unable to direct a stable iOLC identity. In support of this, previous work investigating pericyte to OL reprogramming found that inclusion of *Nkxc.2* in their reprogramming cocktail was refractory to reprogramming (3). In addition to observing the activation of genes unrelated to oligodendrogenesis, genes required for OPC identity were downregulated when cells were transduced with *Nkxc.2* (3).

In a previous study, Khanghahi *et al*. investigated *Sox10-*mediated astrocyte to OLC conversion. Ectopic expression of *Sox10* in astrocytes *in vitro* led to an increase in OPCs at 21DPT (14), a different OLC type and longer time to conversion than we observed in our study. Although similar experimental designs were used in both studies (cortical P3-P5 astrocytes cultured in OPC media in Khanghahi *et al*. versus P0-P5 cortical astrocytes cultured in OPC followed by OL media in our study), these discrepancies may be due to the use of different viral delivery strategies and metrics of reprogramming. In our study, we used lentiviral delivery of *Sox10-*P2A-zsGreen under the control of the long (2178bp) hGFAP promoter (57). In addition, we used Ai14;Aldh1l1-CreERT2 mice and quantified only the iOLCs with an astrocyte origin (based on tdTomato^+^OLC marker^+^expression). In contrast, in the Khanghahi *et al.* study, the authors used a SFFV promoter to deliver *Sox10*-IRES-GFP and reported the number of GFP+ iOLCs in their *Sox10*-transduced cultures. Without an astrocyte specific promoter and stringent lineage tracking, it is difficult to conclude that iOPCs were the result of astrocyte conversion. A non-specific delivery strategy could instead hit a contaminating, perhaps more distantly related cell that would need more time to convert.

In this regard, a recent study suggested that the conversion reported in studies of *in vivo* astrocyte to neuron DLR (6,58,59) was not true reprogramming, but rather the result of erroneous labelling of endogenous neurons due to technical confounds (27). As a result, the DLR community has advocated for the stringent validation of DLR paradigms. In this study, we first examined astrocyte to iOLC conversion in Ai14 cells. This enabled the permanent labeling of transduced cells and therefore, the tracking of those cells through the DLR timecourse. To then validate our findings, we used Aldh1l1-astrocyte fate mapping, which highlighted a lack of conversion with *Nkxc.2*.

When performing our lineage tracing experiments, we also discovered a subpopulation of cells (*Aldh1l1*-tdTomato^neg^zsGreen^+^OLCmarker^+^) that was converted by *Sox10* at a high reprogramming efficiency (Supplemental Figure 5C). scRNA-seq analysis of cells prior to conversion also showed a cluster with low *Aldh1l1* expression compared to the other astrocyte clusters, but with similar levels of *Gfap* expression (Figure 3B), which could explain its transduction by the LV-GFAP::*Sox10*. These *Aldh1l1*^lo^*Gfap*^hi^ cells were predominantly characterized by mature astrocyte markers but also showed expression of genes found in NG2 glia (*Cspg, Pdgfra*) (Figure 3B-C). Curiously, this population did not show expression of proliferation marker *Mkic7* (Figure 3C), in contrast to previous studies showing that astrocyte-like NG2 glia are proliferative (60,61). Of interest, one study reported high co-expression of *Gfap*, *Cspg4* and *Pdgfra* in early postnatal astrocytes (62). This leads to the intriguing idea that these cells represent a population of *Aldh1l1*^lo^expressing astrocytes. If so, *Aldh1l1*-fate-mapping would underestimate reprogramming efficiencies. The presence of these *Aldh11l*^lo^ cells shows that astrocyte cultures are heterogeneous and that different types of astrocytes or astrocyte-like cells may be suitable targets for DLR. Further studies using scRNA barcoding and CellOracle to profile the different types of astrocytes that are amenable to reprogramming will be beneficial in designing more specific and tailored DLR strategies.

Conversion of *Aldh1l1*^pos^ astrocytes to PDGFRa^+^tdTomato^+^zsGreen^+^ iOPCs using *Olig2* occurred at a relatively low rate, with an average conversion efficiency of 16.75% cells. This conversion was even lower for the generation of mature MBP^+^tdTomato^+^zsGreen^+^OLs by *Sox10*, with an average conversion efficiency of 2.83%. This may suggest that although *Sox10* may generate more mature OLCs, it can only do this in a select number of ‘elite’ donor cells. Alternatively, the absence of a substrate to myelinate in our cultures may preclude the true reprogramming ability of *Sox10.* OL survival *in vitro* and *in vivo* has been shown to be dependent on the presence of axons (63). Furthermore, previous studies have shown that mRNA expression of myelin genes is increased when OLs are cultured in the presence of neurons (64), and differentiation of OPCs can be induced with bead or nanofiber scaffolding (65,66). However, additional factors, or the discovery of “better” TFs may also allow for increased reprogramming efficiency. It remains to be determined whether the genes predicted by Fatecode in this study will improve the efficiencies of OLC generation. Nonetheless, it is important to note that the extent of iOLC generation must be balanced with the extent of astrocyte loss as many studies have shown that widespread loss of astrocytes can be detrimental (67,68).

Investigation of the reprogramming trajectory of *Sox10* and Cre-control treated cells showed a bifurcation, where the cells either converted to iOLCs or remained as astrocytes (Figure 3F). It remains to be determined if this is a genetic hurdle or a metabolic hurdle, similar to studies of astrocyte to neuron conversion (69). Future studies investigating the genes that characterize the ‘tipping point’ of conversion will be useful in identifying the mechanisms of astrocyte to iOLC reprogramming. A clearer understanding of how different astrocyte types convert to iOLCs and the mechanisms that underly the reprogramming processes will help us identify the best DLR paradigm for therapeutic applications.

## ACKNOWLEDGMENTS

The authors are grateful to Dr. Lindsey Fiddes at the Microscope Imaging Facility for assistance with live cell imaging. Next-generation sequencing was performed at the Donnelly Sequencing Facility. J.B. was supported by the Baden Havard Endowment Fund, the Ontario Graduate Scholarship and the MITACs Accelerate Fellowship. This work was supported by grants to M.F. from CIHR (PJT-175254), Medicine by Design (Cycle 2 Funding), Temerty Foundation (Pathway Grant), Ontario Institute of Regenerative Medicine (Kickstart Innovation Investment Program), the Stem Cell Network (Early Career Researcher Jump Start Award), the Connaught Foundation (Innovation Grant), CFA-JELF program for microscopy infrastructure, and to S.A.Y. from the CFI-JELF program for scRNA-seq related infrastructure.

## DATA AVAILABILITY

RNA sequence data is deposited in GEO (accession number GSE263185). Genetables are provided as a Cyvers link through the Github page (see methods) housing the scripts used in the manuscript.

## AUTHOR CONTRIBUTIONS

Conceptualization, M.F.; Methodology, J.B., G.D.B., M.F.; Software, E.Y.S., M.S.; Formal Analysis, J.B., E.Y.S., M.S., S.A.Y., M.F.; Investigation, J.B., E.Y.S., A.O., M.S., K.L., M. Fahim, H.T.T., D.L.C., A.D., and S.A.Y.; Writing – Original Draft, J.B. and E.Y.S.; Writing – Review and Editing, M.F.; Visualization, J.B., E.Y.S., M.S.; Supervision, G.D.B., M.F.; Funding Acquisition, S.A.Y., M.F.

## COMPETING INTERESTS

MF, JB are co-inventors on a patent application (Applicants: Justine Bajohr, Maryam Faiz, The governing council of the University of Toronto; Inventors: Maryam Faiz, Justine Bajohr, US Provisional Filed #63/497,357, April 20, 2023).

## SUPPLEMENTARY INFORMATION

**Supplemental Figure 1. Morphological characterization of astrocytes and oligodendrocyte progenitor cells.** Quantification of the **(A)** cell length, **(B)** nucleus size, **(C)** number of branches, and **(D)** branch thickness in Aldh1l1+tdTomato+ and PDGFRα+ OPCs (n= 70 per cell, data are presented as mean ± SEM, each data point represents one individual cell, significance was calculated using Welch’s t-test (A), Mann Whitney test (B,C) and Kruskal Wallis one-way ANOVA (D), ** = p<0.01, **** = p<0.0001)

**Supplemental Figure 2. 14DPT scRNA-seq processing for initial UMAP generation. (A)** Distribution of genes (nFeature), reads (nCount) and percent of mitochondrial reads (percent.mt) per sample. **(B)** UMAP clustering prior to removal of microglia cluster. **(C)**Top upregulated genes characterizing each cluster observed in (B).

**Supplemental Figure 3. Characterization of DLR cultures. (A)** Quantification of contaminating OLCs in post-natal astrocyte cultures 3DPT. **(B)** Quantification of MBP expression in tdTomato^+^ transduced cells 22DPT (n=3, data are presented as mean ± SEM, each data point represents one individual cell culture, a matched pairs one way ANOVA was performed (ns)).

**Supplemental Figure 4. Comparison of DLR clusters 14DPT to established datasets of OLCs. (A)** Violin plots of cell types annotated by Marques *et al.* 2016 within each cluster (OPC = oligodendrocyte progenitor cells, COPs = committed oligodendrocyte progenitors, NFOL = newly formed oligodendrocyte, MFOL = myelin forming oligodendrocyte, mOL = mature oligodendrocyte).**(B)** Violin plots of cell types annotated by Dugas *et al.* 2006 within each cluster (OL = oligodendrocyte, OPC = oligodendrocyte progenitor cells).

**Supplemental Figure 5. Characterization of *Aldh1l1^neg^* and zsGreen^neg^ iOLCs (A)** Quantification of (i) tdTomato^+^ and (ii) zsGreen^+^ PDGFRα^+^iOPCs at 12DPT (n= 3). Data are presented as mean ± SEM, each data point represents one individual cell culture, a paired t-test for (i) and a Wilcoxon test for (ii) was performed. **(B)** Quantification of (i) tdTomato^+^ and (ii) zsGreen^+^ O4^+^iCOPs at 14DPT (n = 4). Data are presented as mean ± SEM, each data point represents one individual cell culture, a paired t-test was used. **(C)** Quantification of (i) tdTomato^+^ (ii) zsGreen^+^ MBP^+^ OLs at 12DPT (n= 3). Data are presented as mean ± SEM, each data point represents one individual cell culture, a matched pairs one way ANOVA with Geisser –Greenhouse correction and Dunnet’s post testing was used (* = p<0.05). **(D)** Representative tdTomato^neg^zsGreen^+^ cell with astrocyte-like morphology at onset (7DPT) and OLC expression at the end (12DPT) of live cell tracking (arrow indicated tracked cell, scale bar = 50um).

**Supplemental Figure 6. DLR timecourse scRNA-seq processing for initial UMAP generation. (A)** Distribution of genes (nFeature), reads (nCount) and percent of mitochondrial reads (percent.mt) per sample. **(B)** UMAP clustering prior to removal of microglia and VLMC clusters. **(C)** Top upregulated genes characterizing each cluster observed in (B).

**Supplemental Figure 7. Differential gene expression of clusters in the Monocle3 trajectory. (A)** Genes enriched in leaf 14 of Monocle3 trajectory. Cells in UMAP colored according to log-normalized gene expression values. **(B)** Genes enriched in leaf 11 of Monocle3 trajectory. Cells in UMAP colored according to log-normalized gene expression values.

**Video S1. Live cell conversion of tdTomato^+^zsGreen^+^ astrocyte to iOLC Video S2. Live cell conversion of tdTomato^neg^zsGreen^+^ astrocyte to iOLC**

**Table S1. Astrocyte and oligodendrocyte progenitor morphology characterization**

**Table S2. 14DPT scRNA seq cell collection metrics**

**Table S3. Pre-transduction, 3 and 8DPT scRNA seq cell collection metrics**

**Table S4. Top 40 genes associated with Fatecode perturbation on node 16 of the latent layer**

**Data S1. BD Rhapsody Whole Transcriptome Analysis whitelist**

**Data S2. Monocle3-derived differentially expressed genes**

## Notes

https://www.ncbi.nlm.nih.gov/geo/query/acc.cgi?acc=GSE263185

https://github.com/eyscott/Bajor_BDRhapsodyData_DLR

### Summary of Updates

Figure axis labels in Figure 2B-D, Supplemental Figure 5A-C and respective figure legends corrected from 'Cre' to 'Control'.

