## Supplemental Figures 1-7, Supplemental Videos 1-2 for "Direct lineage conversion of postnatal mouse cortical astrocytes to oligodendrocyte lineage cells"

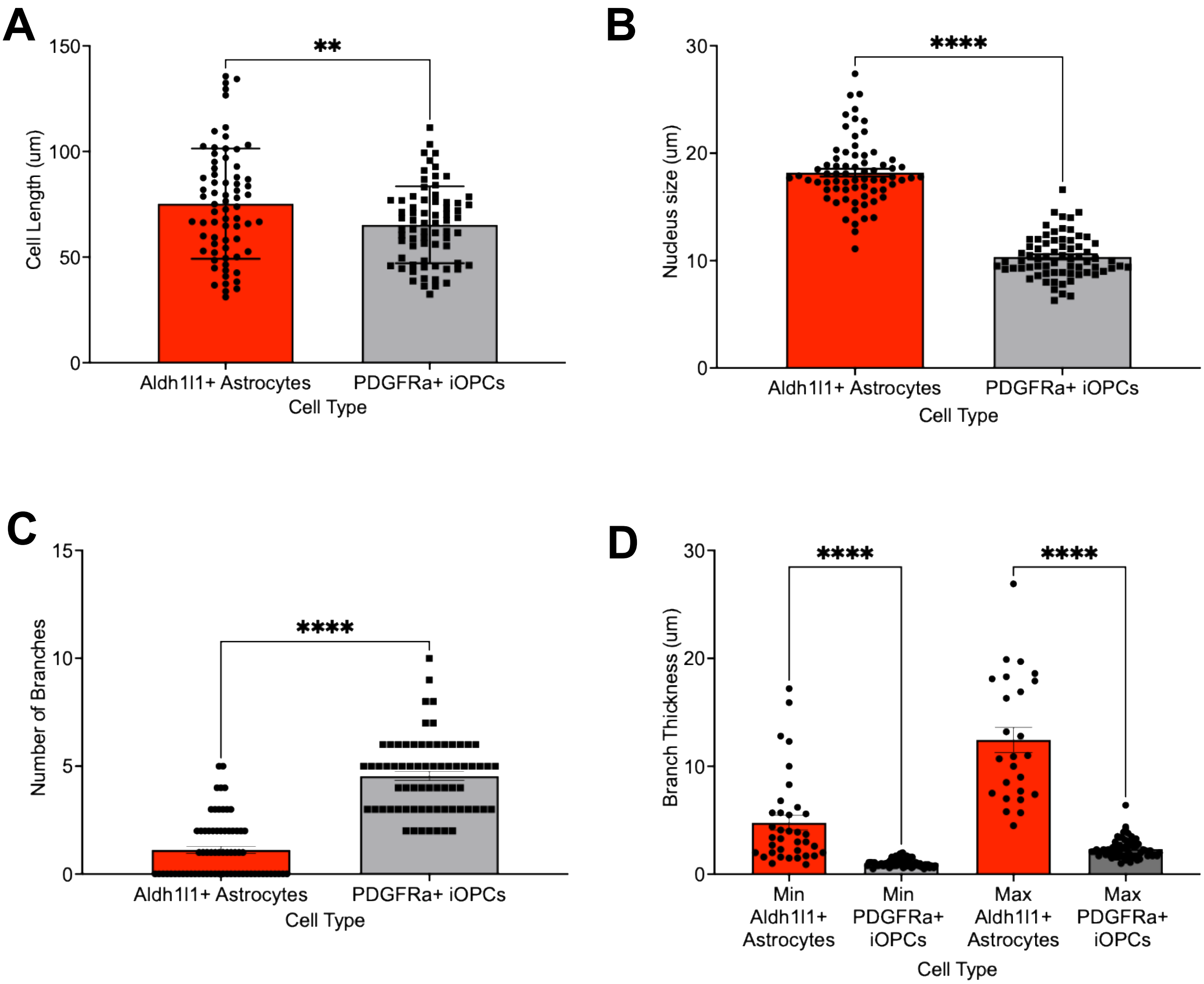
A
Cell Length (um)
150
100
50
0
**
Aldh1l1+ Astrocytes
PDGFRa+ iOPCs
Cell Type
B
Nucleus size (um)
30
20
10
0
****
Aldh1l1+ Astrocytes
PDGFRa+ iOPCs
Cell Type
C
Number of Branches
15
10
5
0
****
Aldh1l1+ Astrocytes
PDGFRa+ iOPCs
Cell Type
D
Branch Thickness (um)
30
20
10
0
****
****
Min Aldh1l1+ Astrocytes
Min PDGFRa+ iOPCs
Max Aldh1l1+ Astrocytes
Max PDGFRa+ iOPCs
Cell Type

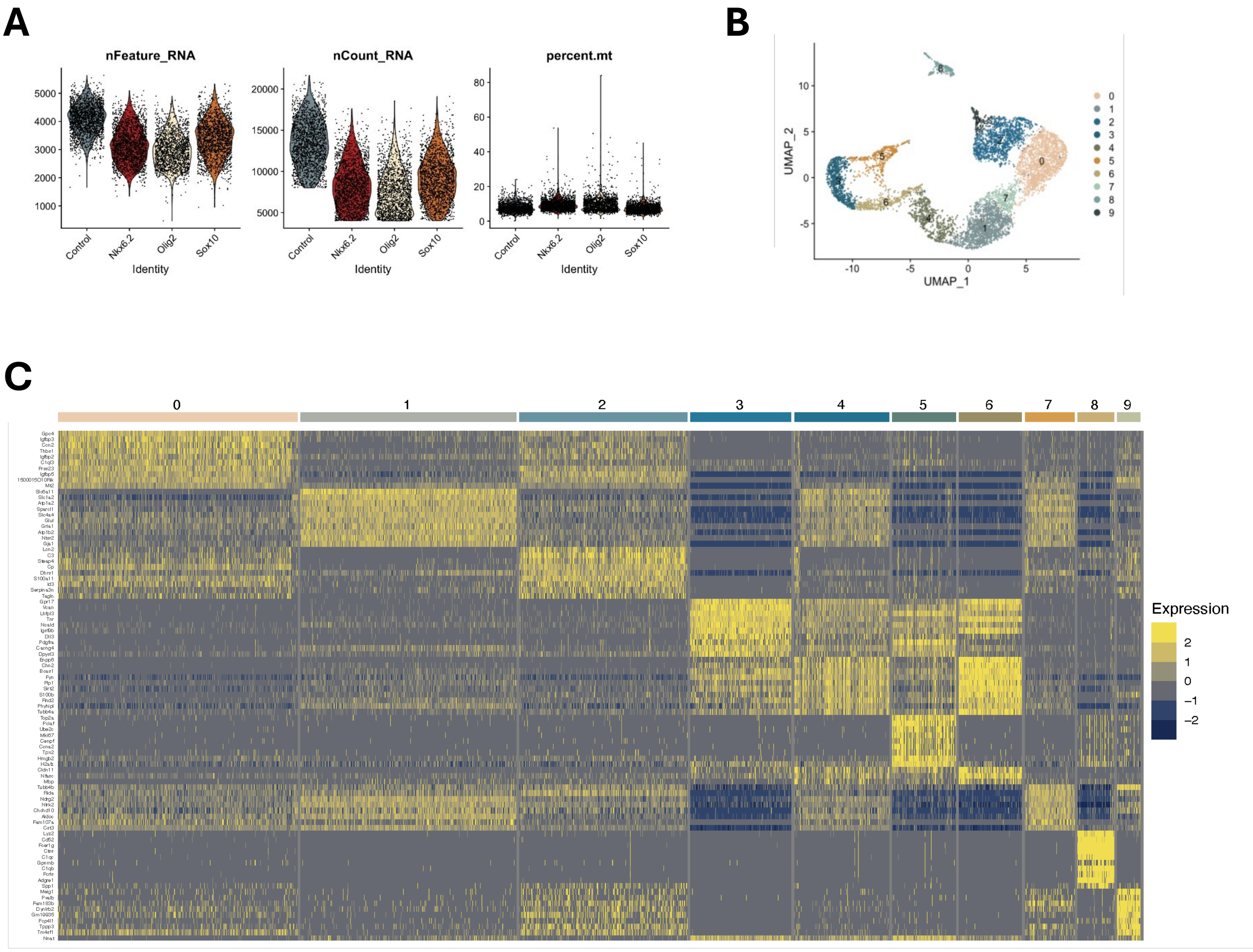
A
nFeature_RNA
nCount_RNA
percent.mt
Control
Nkx6.2
Olig2
Sox10
Identity
B
UMAP_1
UMAP_2
C
Expression

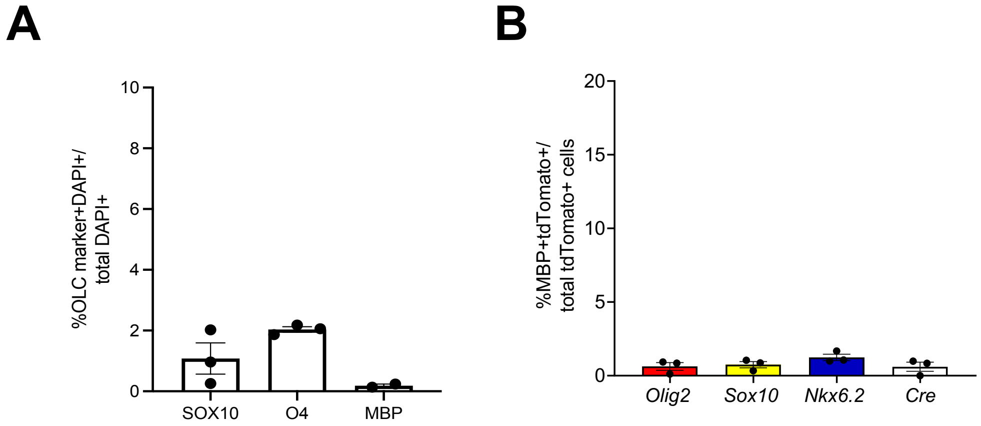

A
%OLC marker+DAPI+/ total DAPI+
10
8
6
4
2
0
SOX10
O4
MBP
B
%MBP+tdTomato+/ total tdTomato+ cells
20
15
10
5
0
Olig2
Sox10
Nkx6.2
Cre

**A**

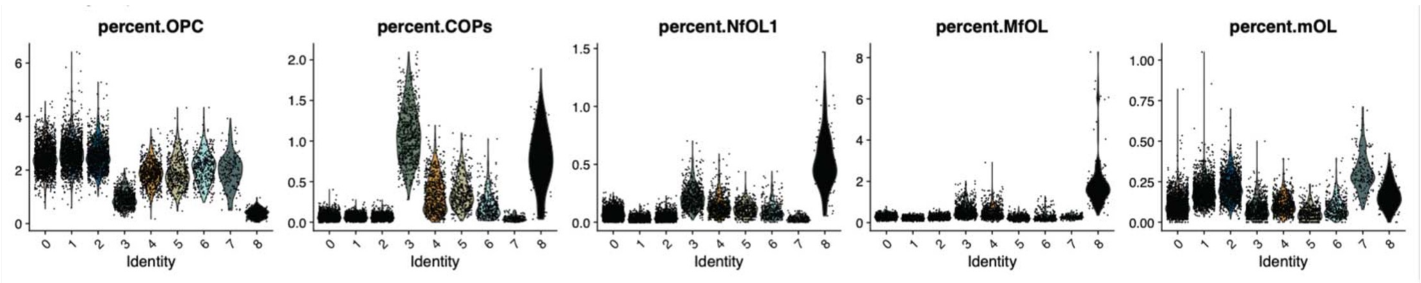
percent.OPC
percent.COPs
percent.NfOL1
percent.MfOL
percent.mOL
Identity
0 1 2 3 4 5 6 7 8

**B**

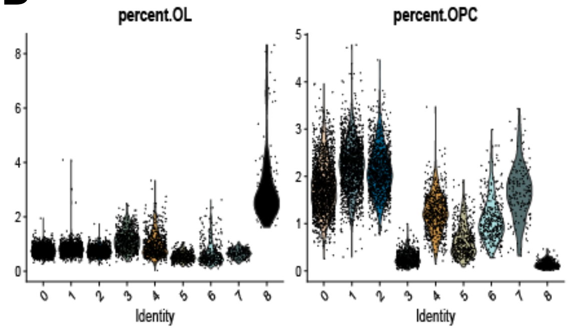
percent.OL
percent.OPC
Identity
0 1 2 3 4 5 6 7 8

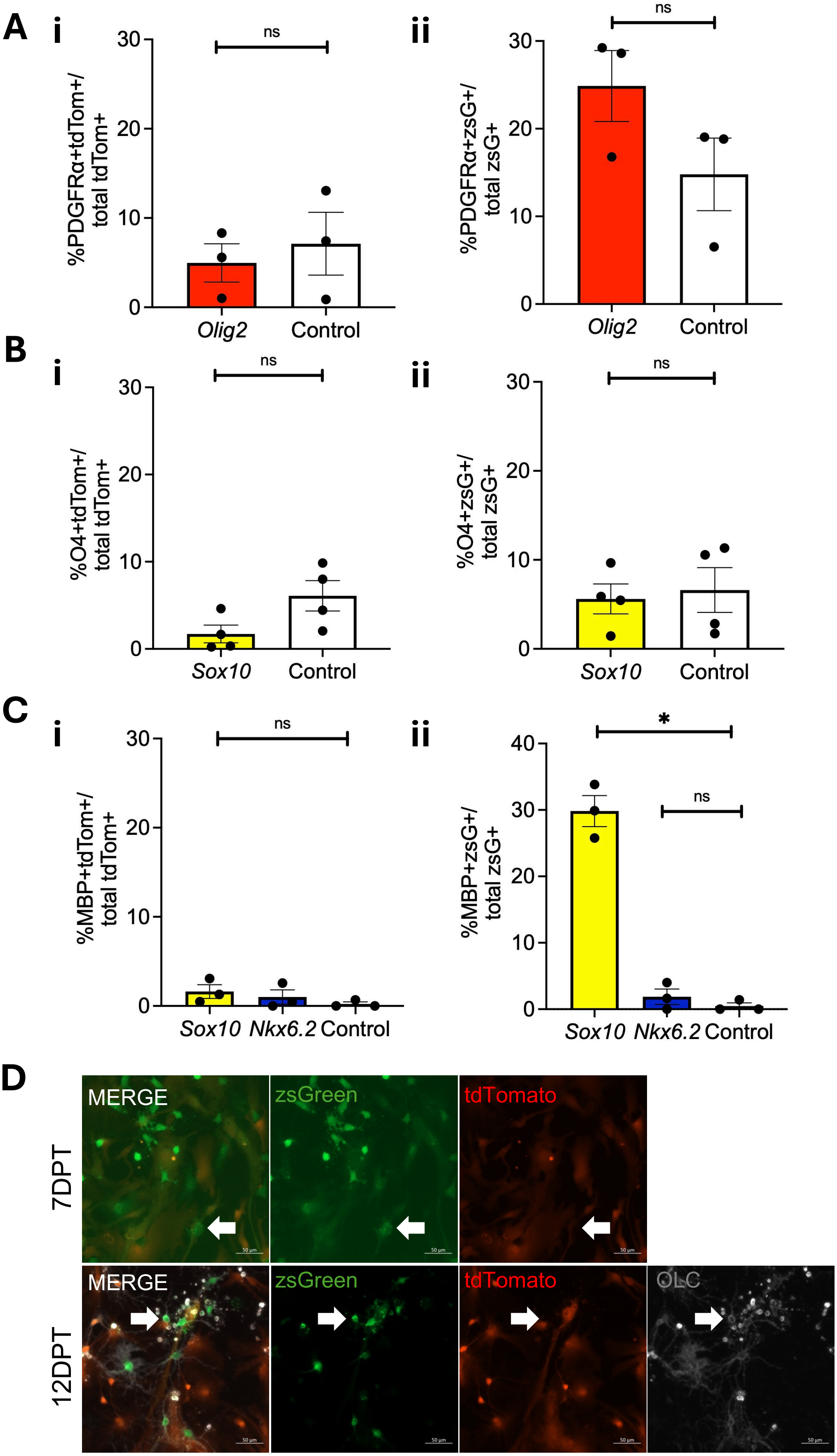

A
i
ii
%PDGFRα+tdTom+/ total tdTom+
%PDGFRα+zsG+/ total zsG+
ns
Olig2
Control
B
%O4+tdTom+/ total tdTom+
%O4+zsG+/ total zsG+
Sox10
C
%MBP+tdTom+/ total tdTom+
%MBP+zsG+/ total zsG+
*
Nkx6.2
D
MERGE
zsGreen
tdTomato
OLC
7DPT
12DPT

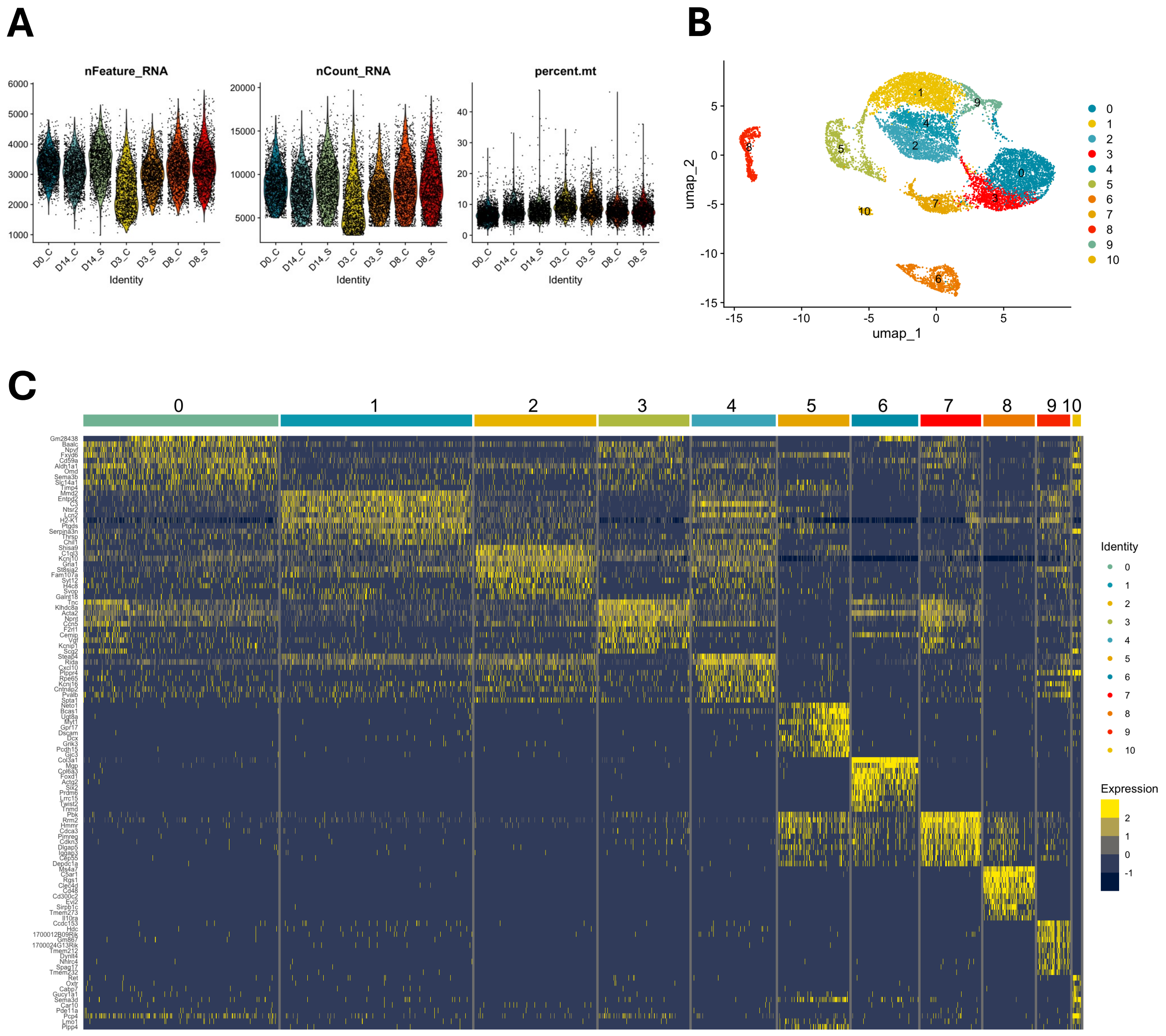
A
B
C
nFeature_RNA
nCount_RNA
percent.mt
Identity
D0_C
D14_C
D14_S
D3_C
D3_S
D8_C
D8_S
umap_1
umap_2
Identity
Expression

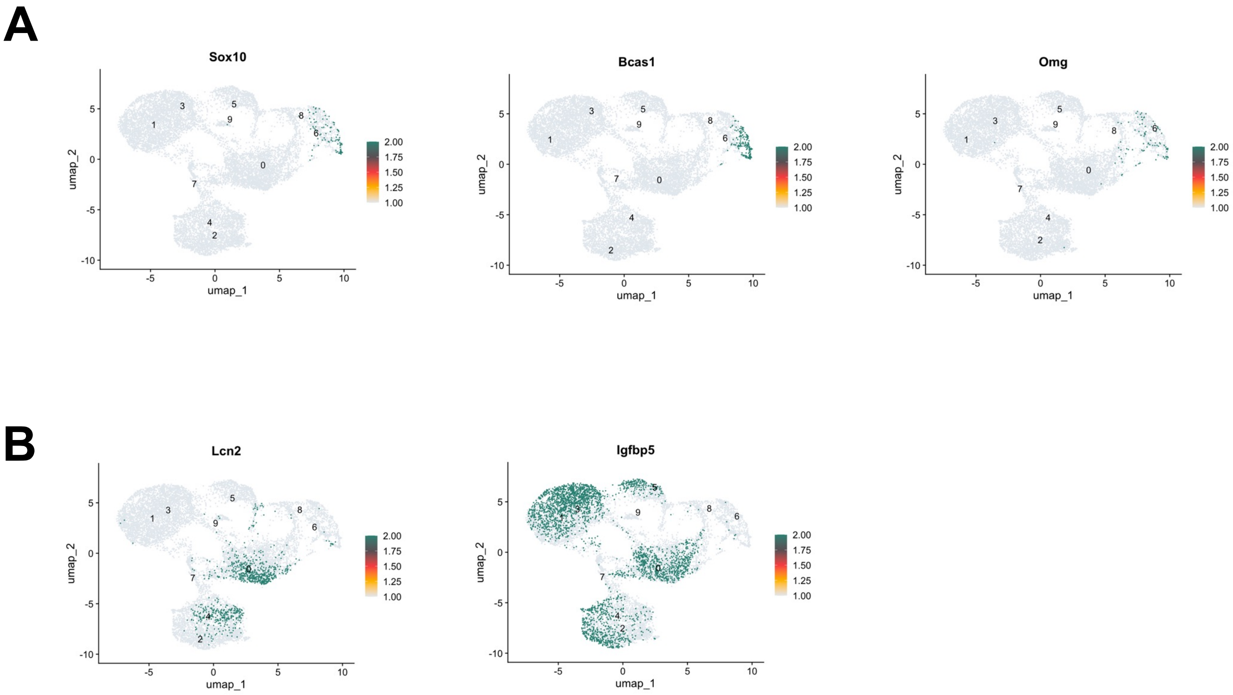
A
Sox10
Bcas1
Omg
B
Lcn2
Igfbp5
umap_1
umap_2
2.00
1.75
1.50
1.25
1.00

**Video S1. Live cell conversion of tdTomato$^+$zsGreen$^+$ astrocyte to iOLC**

https://youtube.com/shorts/GgR_TORW-5s?feature=share

**Video S2. Live cell conversion of tdTomato$^{neg}$zsGreen$^{+}$ astrocyte to iOLC**

https://youtube.com/shorts/FzHlW-SSuBg?feature=share
