## Supplemental Table 2 for "Direct lineage conversion of postnatal mouse cortical astrocytes to oligodendrocyte lineage cells"

**Table S2. 14DPT scRNA-seq cell collection metrics**

| | Control | *Sox10* | *Olig2* | *Nkx6.2* | Average |
|---|---|---|---|---|---|
| **Cell multiplet rate** | 10.1% | 10.1% | 7.0% | 13.3% | 10.13% |
| **Number of viable cells captured in wells at cell load** | 6726 | 12127 | 6733 | 13669 | 9813.75 |
| **Number of wells with viable cells at cell load** | 5967 | 10582 | 6134 | 11546 | 8557.25 |
| **Number of wells with viable cells and a bead** | 4943 | 9322 | 5562 | 8499 | 7081.5 |
