## Supplemental Table 3 for "Direct lineage conversion of postnatal mouse cortical astrocytes to oligodendrocyte lineage cells"

**Table S3. Pre-transduction, 3 and 8DPT scRNA-seq cell collection metrics**

| | **Pre-transduction** | ***Sox10* 3DPT** | **Control 3DPT** | ***Sox10* 8DPT** | **Control 8DPT** | **Average** |
|---|---|---|---|---|---|---|
| **Cell multiplet rate** | 5.7% | 7.0% | 7.7% | 6.9% | 7.0% | 6.86% |
| **Number of viable cells captured in wells at cell load** | 12827 | 10970 | 2750 | 10749 | 14023 | 10263.8 |
| **Number of wells with viable cells at cell load** | 12001 | 9926 | 2480 | 9823 | 12700 | 9386 |
| **Number of wells with viable cells and a bead** | 10172 | 8447 | 2200 | 7824 | 10282 | 7785 |
