## Supplemental Table 4 for "Direct lineage conversion of postnatal mouse cortical astrocytes to oligodendrocyte lineage cells"

**Table S4. Top 40 genes associated with Fatecode perturbation on node 16 of the latent layer**

| Rank | Gene Name |
|---|---|
| 1 | Plp1 |
| 2 | Anks1b |
| 3 | Pllp |
| 4 | Acot1 |
| 5 | Armh4 |
| 6 | Mbp |
| 7 | Tdg |
| 8 | Lrrtm2 |
| 9 | Sox10 |
| 10 | Chd7 |
| 11 | Bzw2 |
| 12 | B3gat2 |
| 13 | Insig1 |
| 14 | S100b |
| 15 | Idh2 |
| 16 | Kcnd2 |
| 17 | Bex2 |
| 18 | Sall3 |
| 19 | Chl1 |
| 20 | Pals2 |
| 21 | Neto1 |
| 22 | Sox12 |
| 23 | Wasf1 |
| 24 | Sox3 |
| 25 | Plppr5 |
| 26 | Spry2 |
| 27 | Fnbp1l |
| 28 | Grm5 |
| 29 | Ppfibp1 |
| 30 | Camsap2 |
| 31 | Ppp1r16b |
| 32 | Ptpre |
| 33 | Serinc5 |
| 34 | B3gat1 |
| 35 | Lrrc17 |
| 36 | Elovl4 |

| | |
|---|---|
| 37 | Mmp15 |
| 38 | Trib2 |
| 39 | Klf9 |
| 40 | Hipk1 |
